# An animal-actuated rotational head-fixation system for 2-photon imaging during 2-d navigation

**DOI:** 10.1101/262543

**Authors:** Jakob Voigts, Mark T. Harnett

## Abstract

Understanding how the biology of the brain gives rise to the computations that drive behavior requires high fidelity, large scale, and subcellular measurements of neural activity. 2-photon microscopy is the primary tool that satisfies these requirements, particularly for measurements during behavior. However, this technique requires rigid head-fixation, constraining the behavioral repertoire of experimental subjects. Increasingly, complex task paradigms are being used to investigate the neural substrates of complex behaviors, including navigation of complex environments, resolving uncertainty between multiple outcomes, integrating unreliable information over time, and/or building internal models of the world. In rodents, planning and decision making processes are often expressed via head and body motion. This produces a significant limitation for head-fixed two-photon imaging. We therefore developed a system that overcomes a major problem of head-fixation: the lack of rotational vestibular input. The system measures rotational strain exerted by mice on the head restraint, which consequently drives a motor, rotating the constraint system and dissipating the strain. This permits mice to rotate their heads in the azimuthal plane with negligible inertia and friction. This stable rotating head-fixation system allows mice to explore physical or virtual 2-D environments. To demonstrate the performance of our system, we conducted 2-photon GCaMP6f imaging in somas and dendrites of pyramidal neurons in mouse retrosplenial cortex. We show that the subcellular resolution of the system’s 2-photon imaging is comparable to that of conventional head-fixed experiments. Additionally, this system allows the attachment of heavy instrumentation to the animal, making it possible to extend the approach to large-scale electrophysiology experiments in the future. Our method enables the use of state-of-the-art imaging techniques while animals perform more complex and naturalistic behaviors than currently possible, with broad potential applications in systems neuroscience.

## Introduction

Many behaviors associated with cognition in rodents are expressed through head and body motion^1^. For instance, foraging^2–4^, olfactory navigation^5,6^, and predator avoidance^7,8^ behaviors are all intrinsically spatial. In addition, head and body motions can correspond to internal deliberation between choices^9–12^. Neuroscience is increasingly exploiting these behaviors to explore the neural mechanisms of cognition by developing tasks based on navigation^13,14^, deliberation between uncertain outcomes^15,16^, integration of information over time^17^, or learning of complex predictive models of their environment^18^. These behaviors have been replicated in head-fixed experiments in simplified form (Fig. 1). However, head-fixation restricts body motion and posture changes, and removes vestibular cues from azimuthal head rotations, which are intrinsically linked to navigation and to many decision making behaviors. The need for head-fixation therefore limits applications of 2-photon microscopy for the study complex cognitive processes.

**Figure 1.**
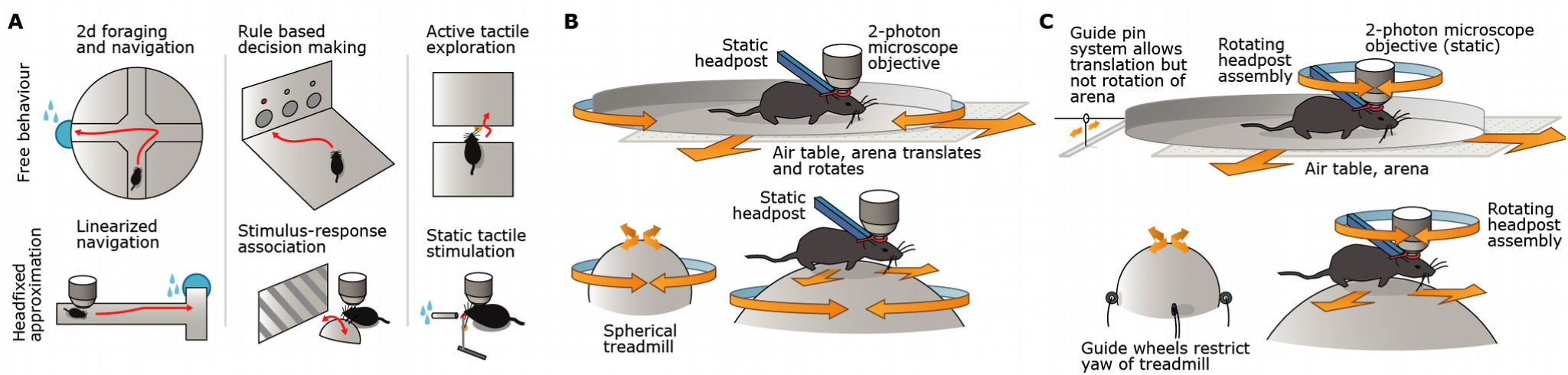
Comparison between free behaviors, virtual reality, and rotating headpost constraint. **A:** Examples of complex decision making tasks in free behavior that are currently restricted in head-fixed VR tasks. i) Complex 2-D navigation with free rotations of the animal, ii) cognitively challenging decision making tasks, and iii) tasks that involve active sensorimotor processes such as vibrissa-based decision making can currently only be replicated in head-fixed and VR environments in simplified form (bottom). **B:** Schematic of conventional head-fixed VR systems that allow 2d-navigation. Mice are head-fixed and the floor they locomote on, either a floating flat arena^25,26^ or a spherical treadmill^45^ rotates under them. **C:** Schematic of head rotation systems. Mice are held in a bearing system that allows horizontal head rotation with low friction and inertia. Animals can therefore rotate their head at their own volition, and can walk on a floating arena or treadmill, which are restricted to translate, without rotation around the vertical axis. These systems fix the animal’s head in the azimuthal plane, but maintain all other degrees of freedom. A static 2-photon microscope, or other recording systems, remains fixed over a brain area of interest.

The predominant approach to circumventing the shortcomings of head-fixed behaviors is to place head-fixed rodents in a virtual reality (VR) environment^19,20^. Animals are placed on a running wheel^21^, disc^22,23^, a floating omnidirectional ball treadmill^24^, or a flat arena on an air cushion^25,26^, and allowed to locomote. Despite the groundbreaking progress of these systems, current VR behaviors lack vestibular inputs, raising questions about whether rodents accept the VR environment as real or whether they merely learn to interact with it^20^. Most saliently, VR systems that do not provide vestibular input suffer from decreased place cell engagement^27^, and decreased selectivity of space-coding neurons^28^. Static head-fixed recordings have only been able to replicate 1-D behavior of grid cells as ‘slices’ through 2-D space^21,29,30^. Cognitive behaviors are similarly limited by head-fixation: the current state of the art are sensory stimulus detection^31–33^, 2-alternative forced-choice or binary discrimination^17,34–36^, and evidence accumulation^37–39^ tasks that often lack the complex decision making and rule-learning observed in free behavior^18,40–43^. In decision making tasks this deficit points to the importance of motor output during deliberation between choices, especially in rodents^10,12,20^. In tactile decision making, this deficit could be caused by the fact that head motion is deeply linked to active tactile sensing trough a number of neural mechanisms^44^ which are disrupted by head-fixation. Current VR and head-fixed tasks are therefore limited in their ability to replicate natural neural spatial coding or to support higher-level cognitive tasks.

Vestibular input can be introduced in head-fixed experiments by rotations of the head^46^ or the entire animal^47,48^, but in most current methods, the rotations are not driven by the animal’s own voluntary motor actions and do not contribute to a sense of agency of the animals; on the contrary, they lead to mismatched signals between the animals’ motor plans, visual, and vestibular input^20^ that can distort the tuning of neurons^49–51^. Natural 2-D grid cell behavior has been observed in VR experiments when rats^45^ and mice^52^ were allowed to rotate under their own volition, suggesting that presence of vestibular input is the key component separating hippocampal and entorhinal function in current head-fixed approaches from free behavior. However, no system with animal-driven head rotations and demonstrated compatibility with 2-photon imaging or attachment of additional instrumentation to the animal currently exists.

A separate approach to address the limitation of head-fixed behaviors is to improve the head-mounted physiological recording systems compatible with free behavior. Progress is being made in miniaturizing recording devices for optical^53,54^, extracellular^55–57^ or intracellular^58–61^ recordings to make them compatible with free behavior. However, the need to miniaturize the devices leads to compromises in data throughput or quality. Head-mounted 1-photon imaging methods impose limits on imaging: no resolution of subcellular compartments, deep tissue access only through implant GRIN lenses, and limited unambiguous separation of individual neurons^54^. Similarly, attempts at miniaturizing 2-photon microscopes^62,63^ have not yet resulted in broadly usable practical solutions. The majority of methods, especially those capable of recording from large populations of neurons, or subcellular activity, require head-fixation. Canonical examples of this are conventional^64–66^ and meso-scale^67^ 2-photon Ca2^+^ imaging^64^ of genetically encoded fluorescent indicators^68^, cell-targeted optogenetic interventions^69–71^, and large-scale extracellular recordings of neural populations^72^, or whole-cell recordings of single^73–75^ or multiple neurons^76–79^.

We have developed a motorized rotating constraint approach that allows mice to freely rotate their heads in the azimuthal plane and locomote in 2-D virtual space of their own volition. The use of a motorized, actively driven rotation system compensates for the weight of the mechanically stable head-fixation system as well as additional recording equipment, and negates the friction of high-stability bearings and slip-rings for signal pass-through. The system recapitulates rotational vestibular input with no residual visual/motor/vestibular mismatch despite head-fixation.

We show that the approach permits the use of high-resolution subcellular 2-photon imaging. Additionally, it allows full use of all recording and intervention methods typically employed in head-fixed animals, as the motor effectively removes the weight of any mouse-attached recording equipment. This system not only facilitates head-fixed studies of navigation, but could also restore a similar sense of agency and spatial decision making to head-fixed animals to recapitulate the complex decision making observed in free behaviors. Our approach can be implemented relatively inexpensively and can be integrated into existing commercial 2-photon microscopes. This new system therefore expands the capabilities of 2-photon imaging to study a wider variety of complex, high-level behaviors.

## Results

The central innovation of our method is the development of an actively force-compensated bearing that allows animals to rotate their heads and bodies freely around one axis, without experiencing significant inertia or friction. This axial constraint system is then combined with a separate system that allows the animal to locomote. Here we use a 2-D air maze^25,26^ that is rotationally constrained (can translate in x and y, but not rotate – see Fig. 9 for details). Through the combination of the 1 degree-of-freedom headpost and the 2 degrees-of-freedom air maze, animals are now free to translate and rotate their heads and bodies, completely removing the visual/motor/vestibular mismatch of current VR systems^20^. The 2-D translational freedom of the air maze and the rotational freedom of the headpost combine to allow free motion-for instance, a head rotation to the left will result in a headpost rotation to the left plus a lateral body translation to the right. Alternatively, the system can be used with a floating-ball treadmill that is equipped with wheels preventing rotation of the ball in the azimuthal plane^45^. The axis of the rotation system is placed over a brain area of interest, which will then remain in the focal spot of an optical imaging system regardless of the animal’s rotation. Similarly, electrophysiological recording equipment or stimulus-delivery systems that are mounted to the rotating frame will remain static with respect to the animal’s head.

The basic function of the system is encapsulated by a simple feedback loop (Fig. 2B). First, a set of strain gages measures the torque applied by the animal to the headpost. Then a controller applies a proportional, but significantly higher torque to the headpost holder via a motor, rotating the headpost as intended by the animal. This applied torque consequently counteracts the apparent torque on the strain gages, and with that, the torque felt by the animal. With a sufficiently high gain, the applied torque overcomes the friction of the bearing, belt drive, and the inertia of the rotating frame. The result is that the headpost holder feels light and virtually frictionless to the animal.

**Figure 2.**
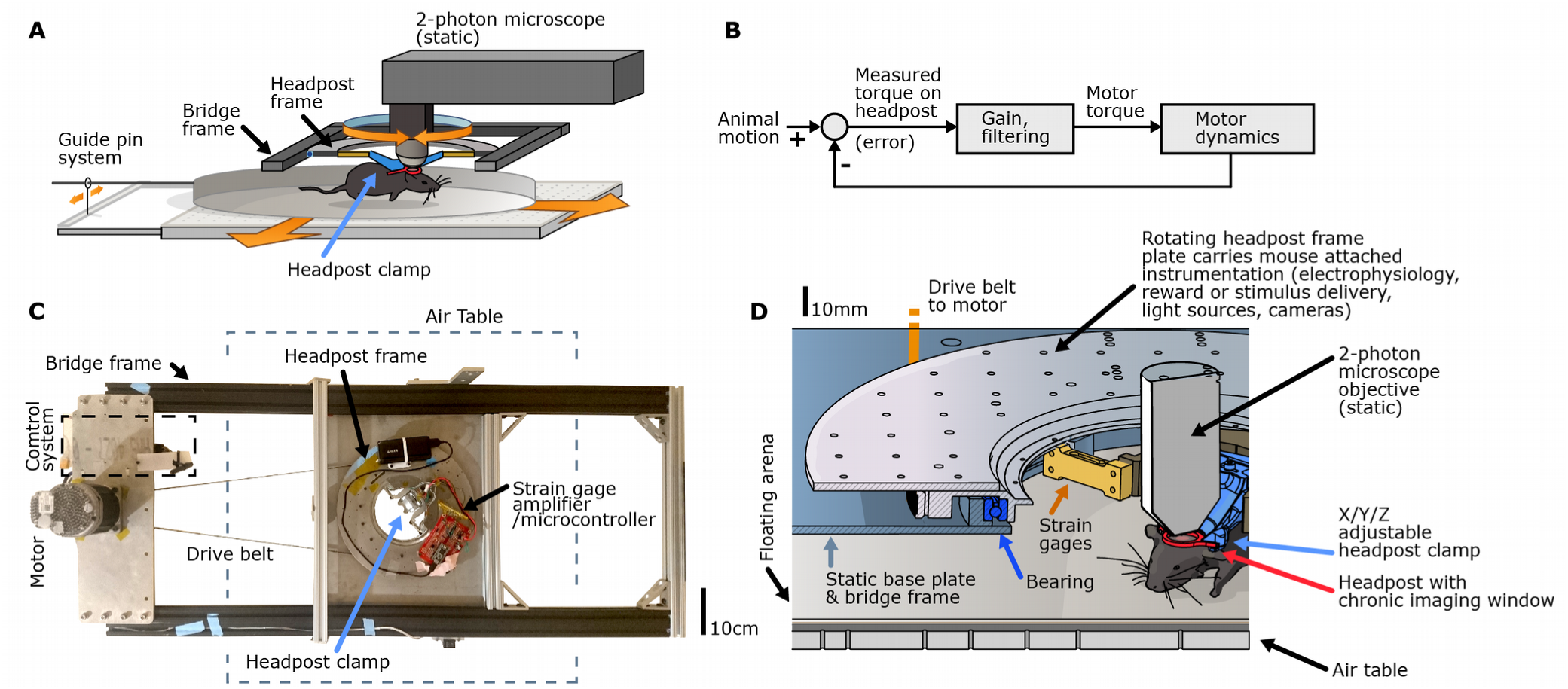
Overview of the system. **A:** Overview of the system integration with a 2-photon microscope. The bridge frame can tilt upward on the air table after the microscope objective is removed, to allow insertion and removal of animals from below. **B:** Basic schematic of the control system. The torque applied by the animal is measured and directly used as an error term by applying a corresponding torque to the motor rotating the headpost frame. **C:** Top-down view of the bridge-frame of the system. **D:** Details of the rotating headpost frame subsystem showing the head post holder that attaches to the rotating frame via a pair of strain gages. The system is made up of a rotating headpost system, a static motor and controller, a translating maze, and an air table.

### Rotational head-fixation maintains basic head-direction encoding

We first verified that a rotating headpost is accepted by mice, and that head-direction coding is maintained in this setting. Intact 2-D grid cells have been observed in head-fixed VR experiments when rats^45^ and most recently in mice^52^ were allowed to rotate under their own volition using a low-friction bearing that restricts head motion to a similar degree as our system. To verify head-direction encoding, we implanted a mouse with a conventional tetrode drive^56^, targeting postsubiculum, where cells encode combined spatial and head-direction direction^80^. We first let mice explore a circular arena with visual cues glued to the walls for ∼15 min while recording spike trains and measuring the mouse’s position with a camera. The mouse’s head direction was coarsely extrapolated from the video data by using motion direction as a proxy. The same mouse was then transferred to the rotating head-fixed system, and a second recording session of 15 min was carried out (Fig. 3B,C). The floating arena for this experiment was similar to that used for free behavior, with comparable visual cues but shorter walls. Spike-sorting was then performed on both sessions together, resulting in a population of identified single units across both settings. We observed re-mapping^81^ of the spatial firing component of the cells, but a good agreement of most head-direction cells between the free and head-fixed settings (Fig. 3D), showing that the head-fixation system does not disrupt the animal’s perception of their head direction.

**Figure 3.**
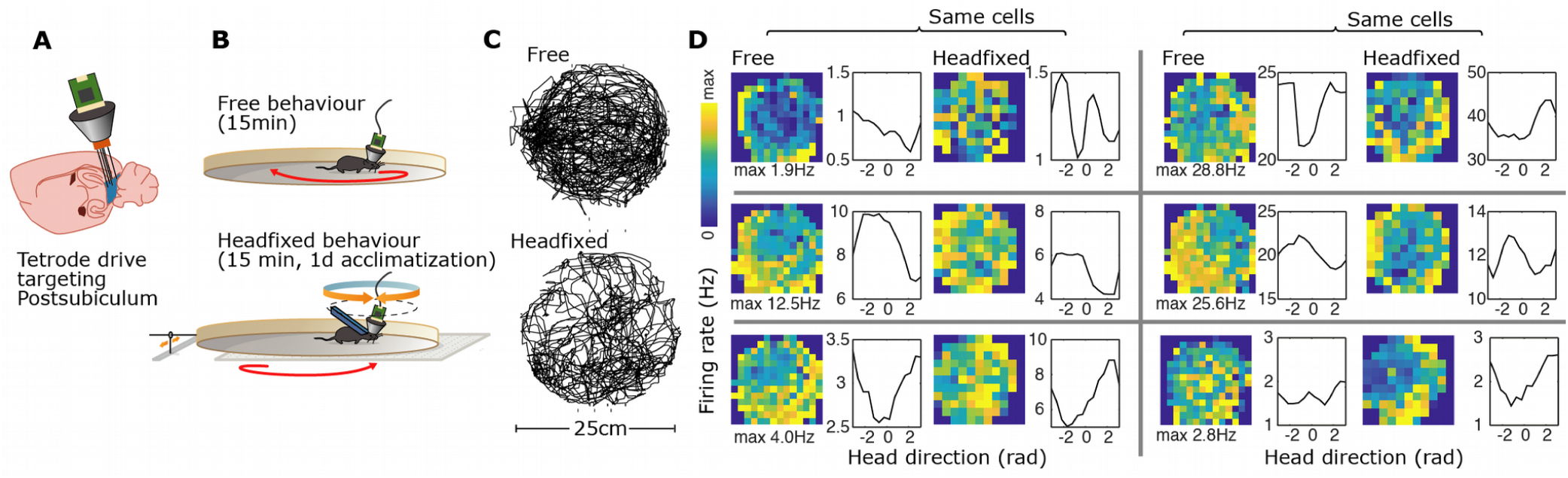
Comparison of head-direction and spatial encoding in postsubiculum across free and head-fixed experiments. **A:** Recording of identified single units in postsubiculum via conventional tetrode drives^56^. **B:** A mouse was recorded during free behavior in a stationary arena while position and head-direction were quantified with conventional videography. The same mouse was then transferred to the head-fixed system and ran for an additional 15 minutes in an arena of the same size and with similar visual landmarks. **C:** After 1 day of prior acclimatization, the mouse explores the arenas both in the freely-moving setting and in the head-fixed system. **D:** Place fields, and head-direction tuning for 6 postsubiculum neurons that show combined spatial and head-direction encoding^80^. The neurons show some spatial re-mapping when the mouse is disconnected from the free-behavior recording system and mounted to the head-fixed system, but a majority of the head-direction tunings are maintained across free behavior and our rotating head-fixation system.

### Rotational head-fixation allows 2-photon imaging of cell bodies and d endrites during locomotion and head-rotation behavior

We then verified the suitability of our method for 2-photon imaging during behavior. We performed 2-photon population imaging of retrosplenial cortex^84^ (RSC) during free foraging. Two mice were implanted with chronic imaging windows over RSC^85,86^ as well as custom headposts, and GCaMP6f^68^ was expressed in RSC neurons using stereotactic injections of AAV.

2-photon imaging in this system poses unique technical issues compared to conventional head-fixed experiments that need to be addressed in order to obtain high-quality data.

First, when using a scanning 2-photon microscope on a rotating sample, the rotation causes image deformations because different lines of the image are scanned at different times, and at different angles (Fig. 4A). We have developed a computational approach for correcting these deformations (See Methods). Second, any deviation of the laser alignment and beam quality from a perfectly axially-centered gaussian beam will cause some unevenness in the image brightness, which poses no issues in static imaging, but causes angle-dependent changes in brightness for ROIs in our method. This artefact can be resolved with a method very similar to the typical baseline fluorescence correction applied to almost all 2-photon calcium imaging experiments^87^ (See methods section, Fig. 10). Finally, the rapid animal motion and rotation afforded by our system cause motion of the brain itself, similar to other settings in which mice are allowed to locomote, requiring a well-calibrated non-rigid image registration method that can remove the residual image deformation. We used the NoRMCorre method^83,88^ for all example data shown here. We verified the suitability of our approach for 2-photon imaging by performing two types of GCaMP imaging experiments that are currently unobtainable with head-mounted microscopes: imaging of dendrite branches and imaging of layer 5 pyramidal somas.

**Figure 4.**
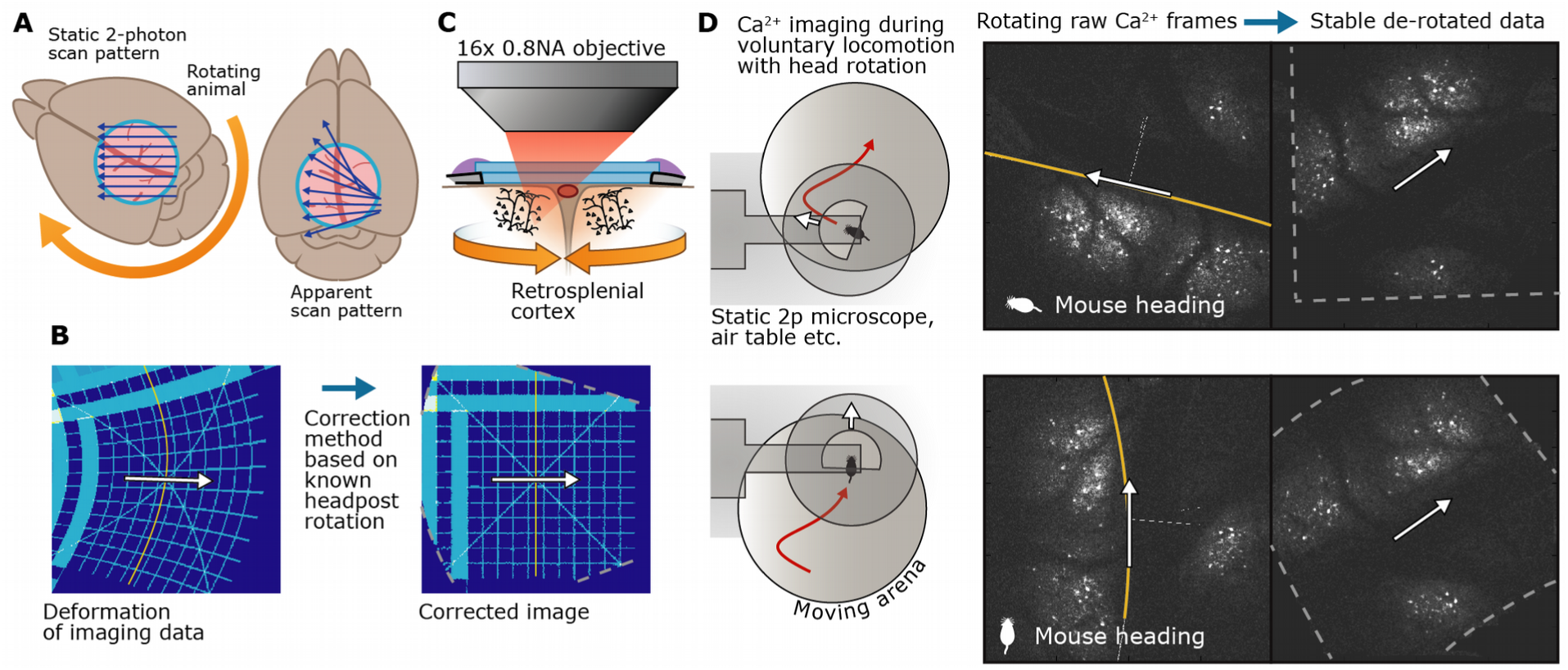
Correction of rotation-induced image deformations. **A:** Schematic of rotation-induced image deformation. **B:** Knowledge of the static scan pattern of the 2-photon microscope and of the animal rotation (known with high precision from the motor encoder) are used to calculate the scan pattern generated by the interaction of the two. The x/y translation for each pixel is then computed from the inferred scan pattern, and a corrected image is generated by interpolating the deformed source image. In moving animals where an additional, unknown x/y motion is present, an additional non-affine registration step^82,83^ is performed in concert with this method. Plots show validation of the correction method on simulated data. **C:** Schematic of 2-photon imaging of populations of RSC neurons during rotation. **D:** Example 2-photon imaging frames from two time points (top vs. bottom) with different head orientations. Left: Schematic of mouse locomotion and rotation in the floating arena. Middle: Raw frames; RSC pyramidal cells in L5 are visible on either side of the central sinus. The raw frames are rotated and deformed by ongoing rotations (indicated by white arrows and image midline). Right: The rotation correction method produces de-rotated images.

**Figure 5.**
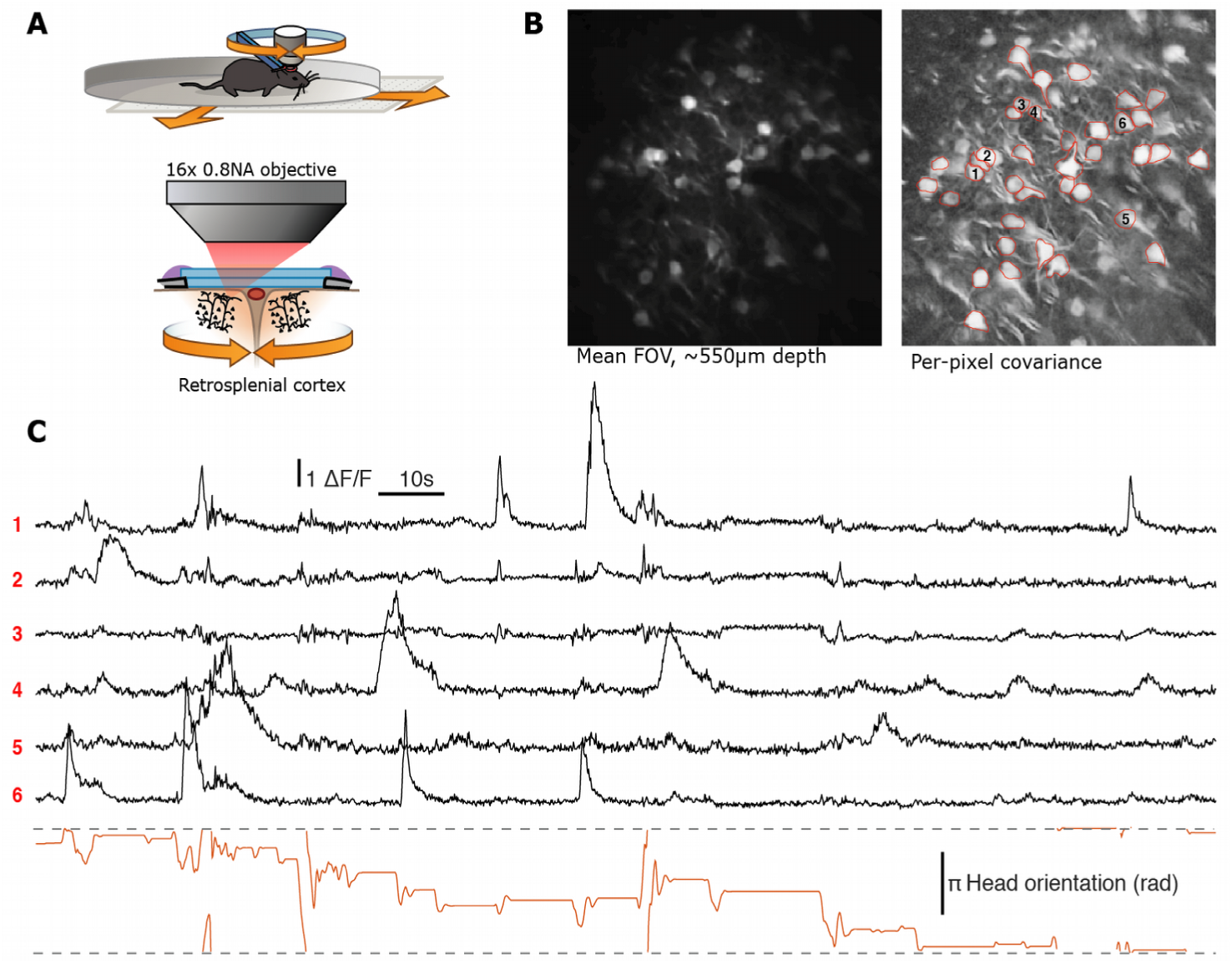
GCaMP6f imaging of cell bodies in cortical layer 5 using the rotating constrain system. **A:** Schematic of the experiment – naive mice were implanted with a chronic imaging window over dorsal midline cortex. We expressed GCaMP6f via AAV injections. Mice were left to explore a circular arena in the rotational headpost restraint during 2-photon imaging of L5 in retrosplenial cortex. Rotated and deformed images were corrected offline prior to analysis (see Methods). **B:** Left, Example field of view from RSC at a depth of ∼550μm below pia (average of 1000 motion corrected frames, see methods for a description of the correction method) showing L5 cell bodies. Right, per-pixel covariance image (covariance of each pixel and its 3×3 pixel environment) and ROI overlays for example cells. **C:** Example ΔF/F traces of cells in A (black), and head-orientation of mouse (orange). See supplementary video for the raw and stabilized videos for this example. No neuropil correction or F_0_ correction other than the correction for rotation was applied to the traces.

Dendrite segments were imaged at ∼100μm (Fig. 6) and ∼50μm (see supplementary movies) below the pia. In both cases, no excessive z motion larger than what we observe in fully head-fixed preparations when mice run on a treadmill was evident, and dendrite segments could be imaged at all head angles. The variation in signal brightness caused by uneven illumination was about an order of magnitude smaller than peak ΔF/F signals and could reliably be corrected by using a simple angle-dependent F_0_ (See Methods).

**Figure 6.**
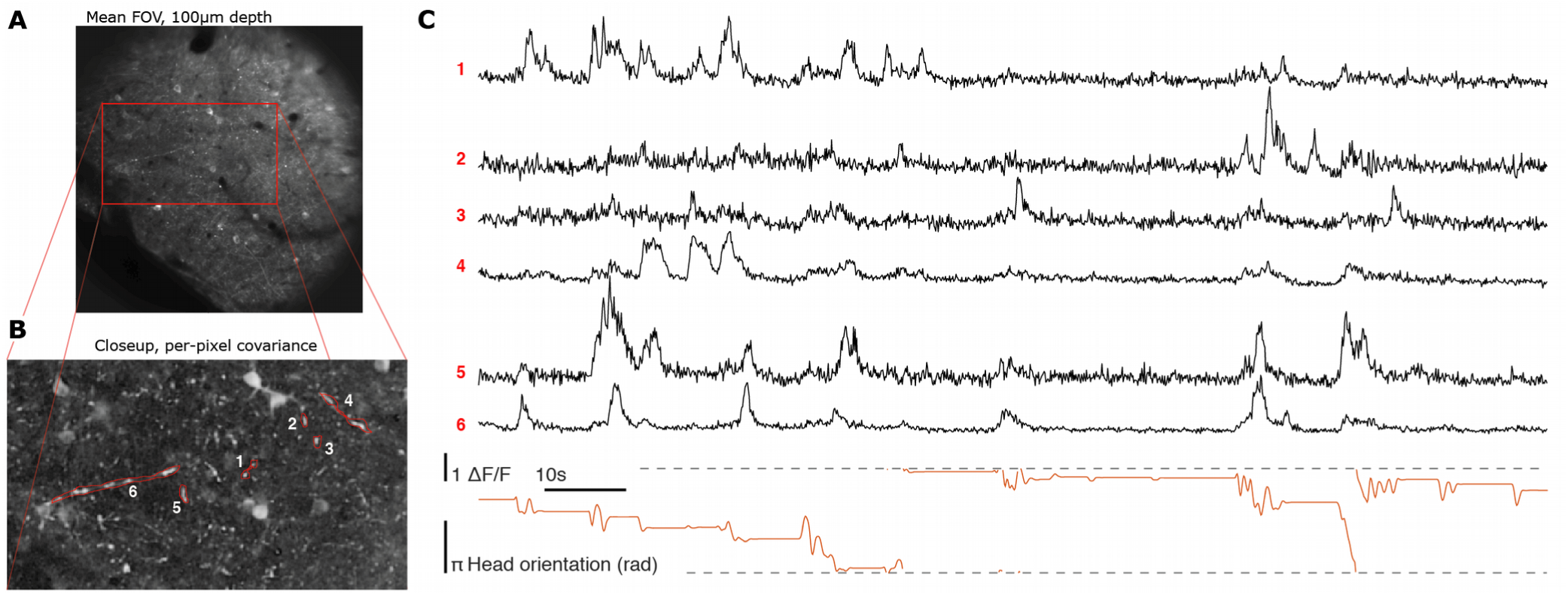
GCaMP6f imaging of dendrite segments in cortical layers 2/3 using the rotating constrain system. **A:** Example field of view from RSC, at a depth of 100μm below pia (average of 1000 motion corrected frames, see methods for description of the correction method) showing a few L2/3 cells and dendrite segments. **B:** Per-pixel covariance image (covariance of each pixel and its 3×3 pixel environment) of a smaller subregion of the FOV. A few example dendrite segments are highlighted. **C:** Example ΔF/F traces of dendrite segments in B (black), and head-orientation of mouse (orange). See supplementary video for the raw and stabilized videos for this example. No neuropil correction, or F_0_ correction other than the correction for rotation was applied to the traces.

We found that appropriate correction algorithms produced imaging quality and stability in the rotational setup comparable to existing head-fixed preparations. Our approach is therefore in principle compatible with any current or future optical methods, including meso-scale imaging^89^ (with modifications of the headpost holder to accommodate larger objectives), deep brain imaging with implanted endoscopes^90,91^ or prisms^92,93^, or single-cell targeted optogenetic manipulations^69–71^.

For brain regions that are further from the midline than RSC, such as somatosensory or auditory cortex, the location of the axis of rotation will not coincide with the midline of the animal. Forward motion of the animals will therefore incur a small torque on their heads. In our experiments such offsets did not appear to have a behavioral effect, but if needed this offset can be compensated by the strain gage amplifier system by re-weighting the measurements of the two strain gages, thereby moving the center-point at which the torque is measured lateral to the rotation axis, to coincide with the animal’s body axis.

### The rotating head-fixation results in small residual torques between the animal and the headpost, and in stable imaging condition despite fast rotations

While motions in the x-y direction can be corrected computationally^82,83^, motion in the z-direction can lead to changes in the brightness of ROIs, or in extreme cases even move entire cells or cell segments in or out of the imaging plane^94,95^.

Z-motion can be caused by two related but separate factors: motion of the headpost itself, due to insufficient stiffness of the heapost holder, or motion of the tissue relative to the headpost and skull due to changes in blood pressure, or strain of the neck and jaw muscles^94^. In extreme cases, such as licking, z-motions of the neural tissue within the skull of up to 15μm have been observed^95^. In order to quantify whether our system is stable under fast rotations, whether it provides enough stiffness to resist z-motion of the headpost, and how it affects motion of the tissue relative to the headpost, we compared recordings of the same ROIs between the rotating headpost, and a control condition in which the headpost was fixed and could not rotate (Fig. 7A). Allowing rotations significantly reduced the torque applied by the animals around the vertical axis on the headpost (Fig. 7B) relative to a fixed headpost (95% CI and medians [0.003, 0.005, 0.036] rotating vs. [0.012, 0.028, 0.162] fixed N=5960 samples, ∼9.9 min, P=<0.0001 ranksum, torque in arbitrary units, Fig. 7E). This result is expected because the rotation system is set up to actively cancel out this torque. In order to quantify the impact of the system on z-motion, we observed the relationship between the torque, and the baseline calcium signal, after removing all activity-driven transients (by analyzing the lowest 10^th^ percentile) on 16 ROIs across the two conditions. If there is z-motion, then torque on the headpost should translate to changes in this baseline fluorescence, as ROIs move in or out of the z-plane. We found that over the range of observed torques (which is smaller for the rotating headpost), the range of baseline fluorescence was [0.76, 0.977, 1.15] in the rotating vs. [0.63, 0.91, 1.25] in the fixed headpost (95% CI and median of 10^th^ percentile of ΔF/F, Fig. 7F). Our torque measurement is not set up to quantify forces other than azimuthal rotational torque, but stability in the z-direction of the headpost itself is not dependent on whether it rotates or not. Our result therefore indicates that the rotating headpost condition can achieve the same, or better stability under animal motion than a static headpost.

**Figure 7.**
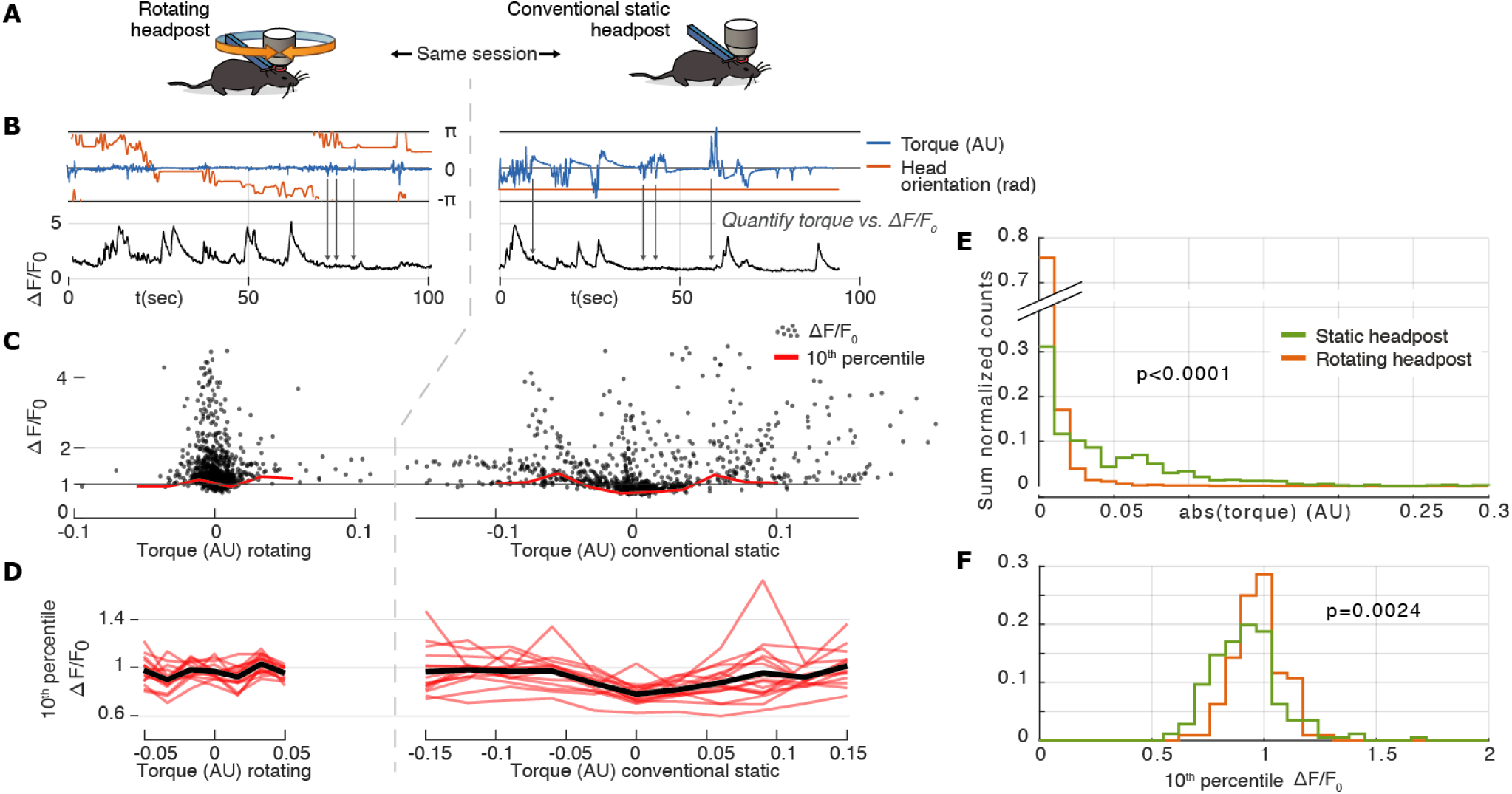
Comparison of headpost forces and imaging stability across rotating and fixed headposts. **A:** Schematic of the experiment: the same ROIs were imaged with the headpost rotation system engaged and during restricted 1-D locomotion (headpost held static, maze floor free to move). **B:** Head orientation and torque measured by the headpost holder across the two conditions. Rotation significantly reduces the torque on the headpost, relative to the conventional fixed headpost. The ΔF/F_0_ for one example ROI is plotted in black. **C:** ΔF/F_0_ of an example ROI plotted against torque across both conditions. Red: 10th percentile of ΔF/F_0_ measures the shift in baseline fluorescence and ignores activity driven transient responses. The dependence of the fluorescent signal on the torque is comparable across the two conditions. **D:** Summary of the dependency of the baseline fluorescence on torque across 16 ROIs. The rotating headpost condition exhibits the same, or better, stability during animal motion than the static case. **E:** Sum normalized histogram of absolute torque values across the two conditions. **F:** Histogram of 10th percentile ΔF/F_0_ values (same as in D).

**Figure 8.**
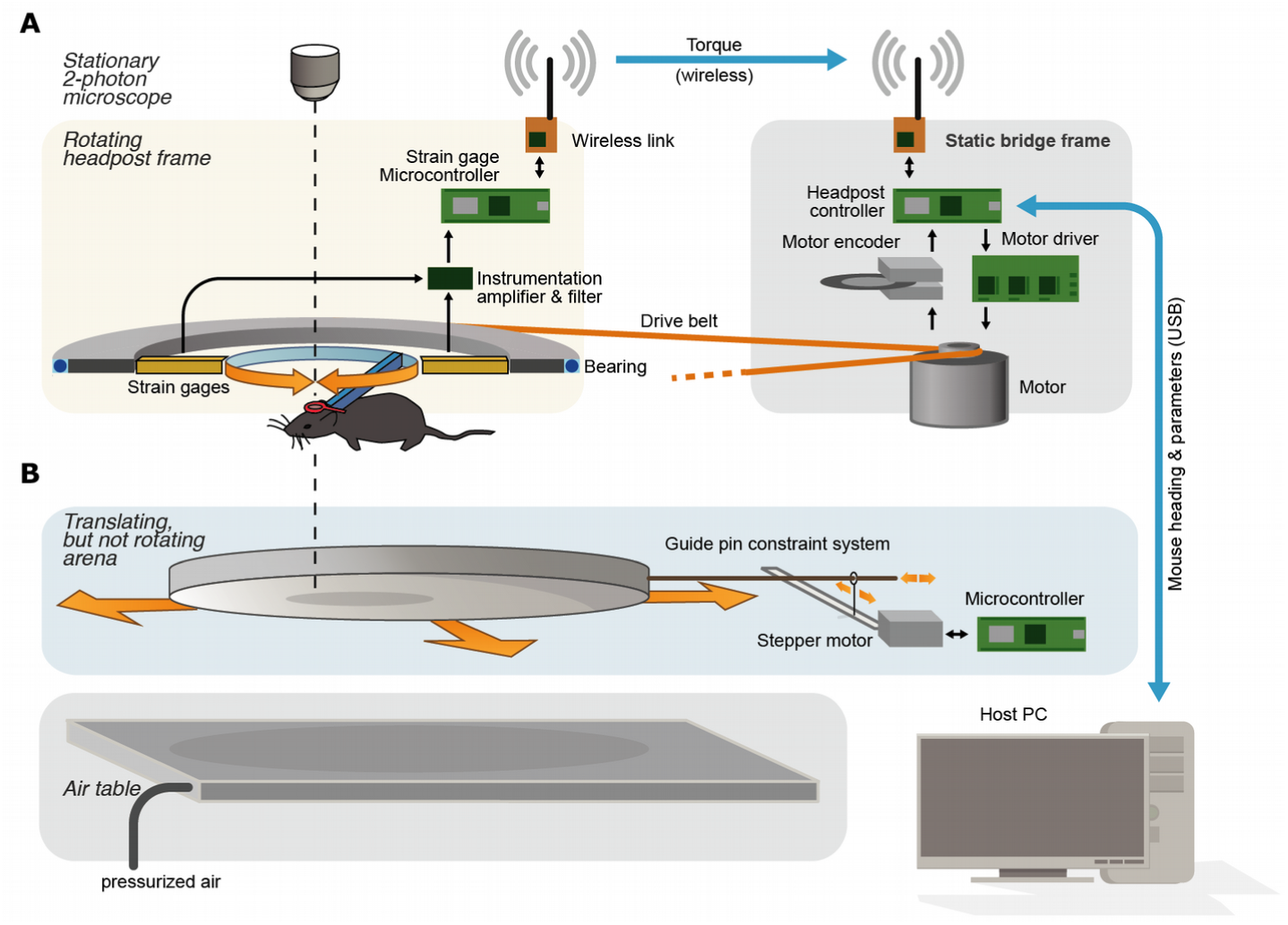
Overview of the subsystems of the method. The system can be divided into two main components, the rotating headpost and the arena. **A:** Headpost system-The *rotating headpost frame* measures the torque applied by the mouse and can hold additional hardware. The static *bridge frame* holds the motor that rotates the headpost frame in accordance with the torque applied by the animal. **B:** Arena system-The *floating arena* moves freely but cannot rotate. The *air table* provides an air cushion for the arena, and tracks the animal’s position within the arena. The arena could be equivalently replaced with a spherical treadmill^45^ or potentially even a flat omnidirectional treadmill^100^.

## Discussion

We developed a relatively simple force-compensated rotating headpost constraint that can be used to achieve volitional head-rotation in behaving mice^45,52^, allowing stable subcellular 2-photon Ca2^+^ imaging (Fig. 5,6). We verified that this approach maintains head-direction coding (Fig. 3), consistent with recent findings of conserved head direction coding and grid cells in VR systems that allow azimuthal head rotation^45,52^.

### Additional data processing steps

Throughout all of our verification experiments, no neuropil correction, time-dependent smoothing or drift compensation via F_0_ was used in order to highlight the image stability of our system. Specifically, we did not apply neuropil correction, a method that subtracts a baseline ‘neuropil’ fluorescence signal extracted from a region around each ROI from its ΔF/F_0_ signal. While useful for correcting for Ca2+ signals from surrounding cells, which are picked up due to the width of the point spread function of the microscope (especially in the z-direction), this method could mask rotation-induced z-motion or image deformations. Further improvements of the signal stability in our system can likely be achieved by applying such methods, given that the brain region studied here, retrosplenial cortex, seems to be characterized by synchronous bouts of widespread activation during locomotion (Fig. 5,6), leading to potentially large contaminant neuropil signal.

### Extensions to whole-cell and large-scale electrophysiological recordings

Many of the same limitations that limit head-fixed behaviors for 2-photon imaging also apply to non-optical recording methods that cannot be miniaturized to the degree required to enable head-mounted free behavior (∼2-3.5 g in mice). Our approach extends to many of these methods because the large load-carrying capacity of the headpost frame, which remains static with respect to the mouses skull makes it possible to attach relatively heavy recording equipment to the mouse’s head. Our current bearing and carrier system supports loads of ∼1-2 kg, but larger drive motors and carriers should be able to scale to larger loads without significant changes to the basic operating principle of the method.

Our approach could therefore facilitate multiple patch-clamp recordings^73,76–78^ and next-generation silicon laminar probes^57,72^ (Janelia/IMEC, Neuropixels consortium). In both cases, the electrophysiology system could be driven from commercially available small form factor desktop computers (Intel NUC or similar) that can mount to the rotating headpost frame, powered by a battery and controlled via WiFi. Alternatively, a slip ring / commutator could be used to route ethernet data to and from the headpost frame. This method removes or simplifies the challenge to route the control signals and data through an electrical commutator.

### Extensions to mouse-attached instrumentation

In the same way that electrophysiological recording equipment can be added to the rotating headpost, other instrumentation can also be attached. We have successfully tested a reward spout using a small solenoid valve, actuated via the same wireless link used for transmitting torque. Similarly, cameras for eye tracking, optogenetic stimulation LEDs, screens for visual stimulation^24^, odor delivery systems^96^, or tactile stimulators^97^ can be mounted. This enables the use of the system in virtual foraging tasks and mixed visual/tactile tasks, as well as sensory control and experimental interventions comparable to those available for classical head-fixed experiments.

### Enabling complex behavior in large arenas

Some behavioral choice tasks can be performed in relatively small behavior chambers^40,43^, while some sensory decision making and navigation tasks require large arenas^98^. We have developed our head-rotation system using a variation of an existing 2-D air ‘maze‘/arena method^25,26^, with the central modification being that our method does not rotate the arena itself. This significantly reduces the problems caused by the inertia of the rapidly spinning arena in current approaches^25,26^, which will make it possible to use large arenas. Our prototype system uses a 25cm diameter floor plate, but diameters of up to 50cm should be possible. Additionally, because the system rotates the animal, and the arena only translates, it will be practically feasible to introduce reward ports and/or stimulus delivery systems. Lightweight reward ports can be attached to the arena and water/odorant/food delivery tubes can be left hanging from the arena to an overhead support, allowing motion of the arena. Similarly, tactile, visual, or odorant stimuli can be delivered through openings in the arena walls that the mouse can extend its head through, with the stimulus delivery system moved into position via a linear actuator. A similar system has been successfully integrated into a conventional air-maze system^26^. Our approach also allows the use of actively-actuated 2-D treadmills^99,100^, or of ball systems that are actively (via motors^101^) or passively (by rollers^24,45^) restrained from rotating around the vertical axis. The system also supports scaled-up headposts on a larger arena or treadmill for use in rats^45^ or other rodents such as tree shrews^102–105^. The downside of spherical treadmill systems is that the styrofoam balls provide less natural posture and locomotion than the flat arena and lack tactile feedback such as walls and additional maze elements^26,89^. The upside of styrofoam ball systems over the flat arenas is that the virtual environment can be made arbitrarily large^19,24^.

In sum, we have developed an approach that allows mice to freely rotate their heads in the azimuthal plane and move in 2-D space under their own volition, maintaining rotational vestibular cues during 2-photon imaging. This approach will enable head-fixed studies of navigation, as well as other behaviors that are enabled or facilitated by allowing the animals to rotate their heads, that make use of 2-photon imaging and other recording methods that require head-fixation. This system therefore provides the ability to image using state-of-the-art imaging techniques as animals perform complex and naturalistic behaviors, with broad potential applications in systems neuroscience.

## Methods

### Mechanical system

#### Headpost clamp

We used a slightly adapted standard headpost and clamp^85,93^. The headpost clamp is engineered so it can be tightened or released with a single thumbscrew allowing easy insertion or removal of mice with one hand.

#### Strain gage frame

An aluminum frame connects the headpost clamp to an array of 2 strain gages (Phidgets Inc. load cells, See Fig. 2) that are set up as an opposing pair to not register any forces other than torque around the rotation axis. The torque is measured via a battery powered instrumentation amplifier and transmitted via a wireless link (SparkFun RFM69) to the main micro-controller (Teensy 3.6).

#### Rotating frame

The strain gages are held in a rotating frame that is made from milled aluminum on a flat waterjet cut base plate (1/4” aluminum). This rotating frame also holds all rotating amplifiers, control electronics, wireless link, and a battery system. A rechargeable lithium ion external cell phone battery with ∼5000mAh is used to power the microcontroller and amplifiers. The frame also includes a bearing carrier surface for a thin profile bearing (Kaydon K13008XNOK), and a sprocket ring for the drive belt.

#### Air maze system

The maze that the animals walk on needs to be translationally friction free and have low inertia ^25,26^, but cannot be allowed to rotate. The air maze itself is a circular, or otherwise shaped arena of ∼25 cm diameter, that floats on an air cushion providing animals with the ability to walk without experiencing resistance. The air maze floor is made from carbon fiber sheet. Crucially, the air maze needs to be restricted from spinning on its own, so that all torques exerted by the animals are transferred to the headpost and can be compensated there. For this purpose, the arena has a protruding constraint pin that fits through a corresponding guide (precisely, a pair of guides, see Fig. 9B,C) on the constraint system shuttle, acting as a linear bearing: The guide pin can slide in and out of this bearing, allowing the arena to move in the x direction (Fig. 9C). In order to allow y-direction motion, the system measures the torque or angle with which the guide pin enters the bearing, and actively keeps the angle between the constraint shuttle and the guide pin at 90 degrees. The shuttle therefore constantly follows the arena in the y direction if the arena translated, but rotational torque does not move the shuttle and is absorbed by the rigid constraint system (Fig. 9D). Because the air maze does not rotate, electricity and signal wires can easily be routed through the guide pin, or by slack lightweight wires or tubes, and need no commutation. The motion of the arena is tracked with an IR LED driven by an adjustable current source^106^ that is attached to the arena and is tracked with a pixy camera^107^. The air maze floats on an air cushion provided by a large air table made from two 30 x 30” sheets of clear acrylic, the top of which has a pattern of small holes drilled. The space between the acrylic sheets is pressurized with air from building utility air. A series of threaded rods tie the two sheets together to ensure that the distance between the acrylic plates stays constant despite the air pressure.

**Figure 9.**
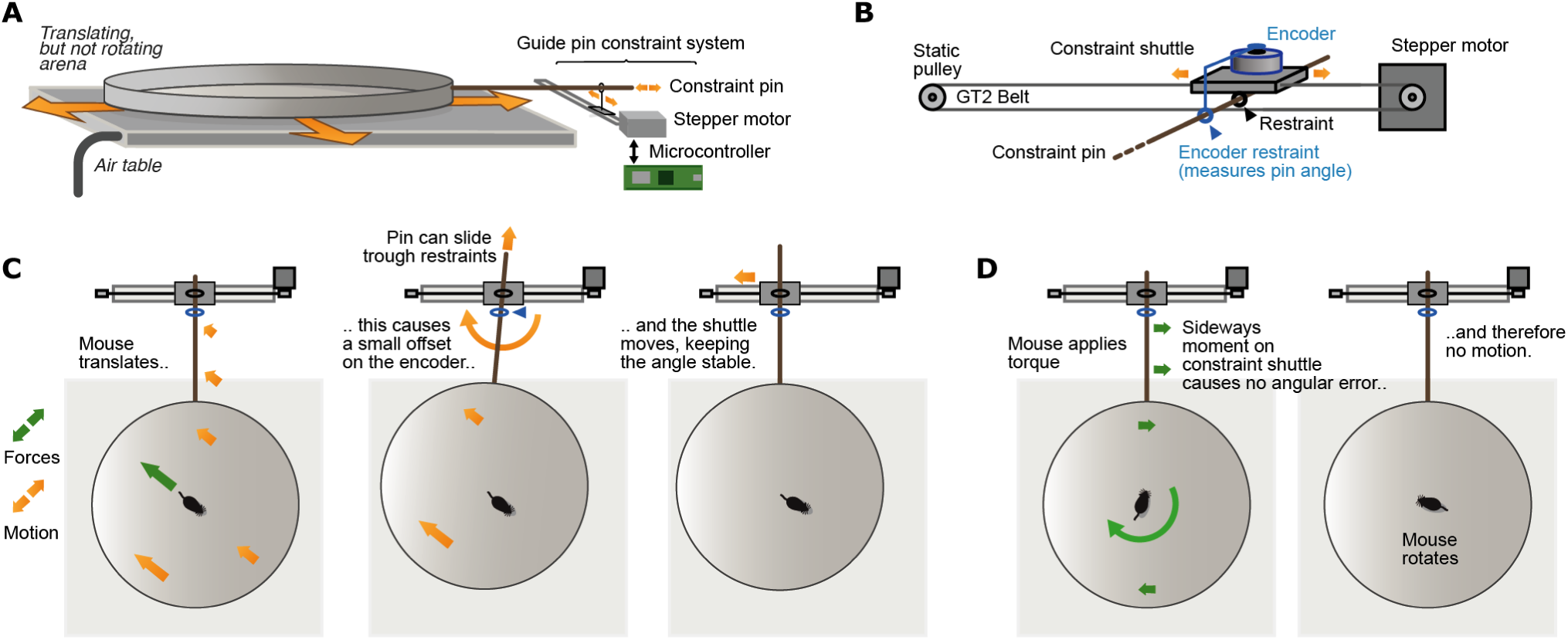
Guide pin system for restricting maze rotations but allowing translations. **A:** Schematic of the system. A guide pin is rigidly attached to the floating maze^25,26^, and prevented from changing its angle by the constraint system. The constraint system is attached at one side of the air table, the width of the system and the length of the guide pin are equal or larger than the desired x/y motion range, or the diameter of the maze. **B:** The constraint system is made up of a shuttle with two restraint rings (which act as linear bearings and allow the guide pin to slide in or out freely). The two restraints measure the angle at which the guide pin enters the shuttle, by means of an encoder (a strain gage can also be used here). The whole constraint shuttle can be moved laterally via a stepper motor on a toothed belt. The belt tension itself is sufficient to hold the shuttle in place, but an additional linear rail can be used if needed. **C:** When the mouse moves, the restraint system measures the resulting small angle change and stops the maze from rotating by keeping the restraint shuttle at the same angle to the maze. **D:** When the mouse applies torque instead of translational motion, the guide pin does not change the angle by which it enters the constraint shuttle. This causes no motion of the shuttle, which therefore resists the torque applied by the mouse.

#### Bridge frame and motor

The static bridge frame holds a bearing carrier surface for the main bearing, the base plate that forms the non-moving ‘ceiling’ that is seen by the animals, and can be covered with a mirrored surface or fitted with screens for display of a VR environment. The bridge frame also carries the drive motor and motor control electronics. The whole bridge frame can be tilted upwards, allowing easy insertion and removal of animals. The main microcontroller receives torque data from the rotating frame system at ∼500 Hz and computes the required motor torque. The rotating frame is driven via a Gates GT2 belt from a brushless motor (Teknic Clearpath) mounted on one side of the bridge frame. The motor is controlled by a separate microcontroller via a simple PWM torque command. The microcontroller also reads the position of the motor encoder in order to track the heading of the mouse. The controller then sends this heading to the host PC via a stream of serial data.

### Surgery

Mice (C57BL/6) were aged 8-15 weeks at the time of surgery. Animals were individually housed and maintained on a 12-h cycle. All experiments were conducted in accordance with the National Institutes of Health guidelines and with the approval of the Committee on Animal Care at the Massachusetts Institute of Technology (MIT). All surgeries were performed under aseptic conditions under stereotaxic guidance. Mice were anesthetized with isofluorane (2% induction, 0.75–1.25% maintenance in 1 l/min oxygen) and secured in a stereotaxic apparatus. A heating pad was used to maintain body temperature, additional heating was provided until fully recovered. The scalp was shaved, wiped with hair-removal cream and cleaned with iodine solution and alcohol. After intraperitoneal (IP) injection of dexamethasone (4 mg/kg), Carprofen (5mg/kg), subcutaneous injection of slow-release Buprenorphine (0.5 mg/kg), and local application of Lidocaine, the skull was exposed. For some mice, AAV was injected as described. The skull was cleaned with ethanol, and a base of adhesive luting cement (C&B Metabond) was applied.

Chronic headpost and imaging window implants were performed over central midline cortex. 4 to 6 injections of ∼50nl each of AAV2/1-hSyn-GCaMP6f (HHMI/Janelia Farm, GENIE Project; ∼1012 viral molecules per ml)^87^, were made bilaterally, around 0.3-0.6mm from the midline at depths of ∼500μm. Mice were given 2 weeks to recover and for virus expression before the start of recordings. A 3 mm craniotomy was drilled, virus was injected, and a cranial window^85,108^ ‘plug’ was made by stacking two 3 mm coverslips (Deckgläser, #0 thickness (∼0.1 mm); Warner; CS-3R) under a 5mm coverslip (Warner; CS-5R), using optical adhesive (Norland Optical #71). The plug was inserted into the craniotomy and the edges of the larger glass were sealed with Vetbond (3M) and cemented in place. The dura was left intact.

Chronic drive implants were performed identically to the window implants, but instead of a 3mm craniotomy for a glass window, a 2mm craniotomy was drilled ∼2mm lateral and ∼0.5mm anterior of the transverse sinus. A durotomy was performed, and tetrode drives^56^ were implanted and fixated with dental cement.

### Data processing

All data were acquired at a frame rate of 9-11Hz using a 2-photon microscope with a combined resonant scanner/galvo system (Neurolabware, Los Angeles, CA). Imaging was performed with an excitation wavelength of 980nm. The rotation of the animal was read out from the drive motor’s encoder at a frequency of 500Hz and was saved together with the imaging data.

#### De-rotation

The imaging data recorded while the animal rotates is itself rotated relative to the scan system of the microscope, and therefore distorted. The distortion results from the fact that the image acquisition is not instantaneous, so that the top line of a frame will be acquired at a different time, and possibly at a different angle of rotation, from the bottom line (Fig. 4A). In mild cases this results in an apparent curving of the image, theoretically it can lead to parts of the field of view getting imaged twice per frame. We have developed a computational method for correcting these distortions. The method works by computing the ‘forward’ model of the distortion by generating a map of the x and y distortions per pixel by recapitulating the scan using the true scan speed and the measured animal rotation for each image line. The resulting distorted x/y positions of each original x/y pixel are then used to reverse the deformation (Fig. 4B-D). After this correction step, images are processed with a standard motion correction pipeline^82,83,109^. After motion correction, ROIs of individual cells and dendrite segments are identified manually using custom software and ΔF/F_0_ traces are computed.

#### Correction of uneven illumination

If the illumination of the FOV is not completely even, due to small deviations of the laser alignment from the objective or rotation axes, or non-uniform back-aperture illumination of the objective, rotation of the animal will bring imaged structures in and out of areas of higher or lower brightness. In our data, this effect accounted for only ∼10-20% of ROI brightness for ROIs near the center of the field of view, approximately the center 50% of max FOV. To resolve this artefact we adapted the method typically used for correction of baseline fluorescence^87^: Typically, for each ROI, a baseline fluorescence F_0_ is computed as the 2-15^th^ percentile of either all fluorescence data for a session, a pre-stimulus period, or in a moving window of a minute or more. This F_0_ is then used as a baseline and the time series ΔF/F_0_ is used for subsequent analyses. Here, we used a similar method, but F_0_ is computed not over time, but across angles (Fig. 10B) by computing a quantile for each angle. On our experiments, this simple approach sufficiently corrected the angle dependent brightness changes (Fig. 10B,C). No neuropil correction or F_0_ correction other than this static correction for rotation was applied to the traces.

**Figure 10.**
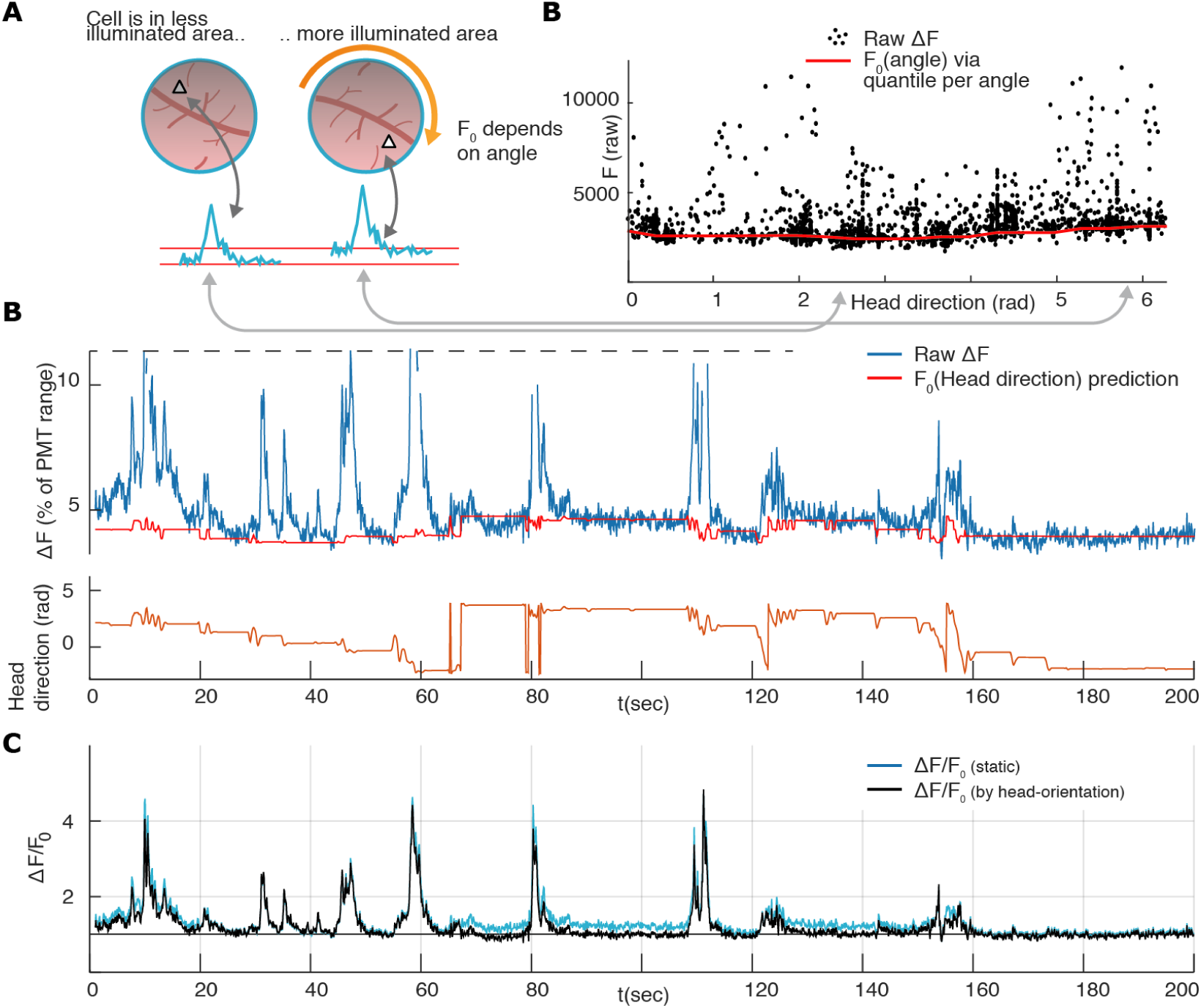
Correction of angle-dependent brightness changes. **A:** Uneven image brightness leads to angle-dependent changes in ROI brightness. **B:** Method for correcting these angle-dependent brightness changes. For each ROI and for each angle, a F_0_ is estimated by computing the 20^th^ percentile of the ROI fluorescence. This method is analogous to the standard method for computing F_0_, but is applied over angles rather than over time. **B:** The resulting lookup-table predicts F_0_ from the current angle. The example trace shows that the resulting F_0_ predictions tracks the baseline fluorescence well, and that the brightness modulation over the full range of rotations is well below the amplitude of the calcium transients. **C:** Resulting ΔF/F_0_ for the example ROI (black) and same trace when a static F_0_ is used (blue). In this example, no additional time-varying F_0_ is used.

## Acknowledgements

We thank Jonathan P. Newman for many helpful discussions, technical advice and support. We also thank Matt Wilson for helpful discussions of the method. We thank Lou Beaulieu-Laroche, Lukas Fischer, Marie-Sophie van der Goes and Enrique Toloza for comments on the manuscript. Funding was provided by the MIT Research Support Committee -NEC Corporation Fund for Research in Computers and Communications (M.T.H.), and the Simons Center for the Social Brain at MIT postdoctoral fellowship (J.V.).

## References

1. Kawai, R. et al. Motor Cortex Is Required for Learning but Not for Executing a Motor Skill. Neuron 86, 800–812 (2015).

2. Kolling, N., Behrens, T. E. J., Mars, R. B. & Rushworth, M. F. S. Neural Mechanisms of Foraging. Science 336, 95–98 (2012).

3. Morris, D. W. & Davidson, D. L. Optimally Foraging Mice Match Patch Use with Habitat Differences in Fitness. Ecology 81, 2061–2066 (2000).

4. Stopka, P. & Macdonald, D. W. Way-marking behaviour: an aid to spatial navigation in the wood mouse (Apodemus sylvaticus). BMC Ecol. 3, 3 (2003).

5. Eichenbauma, H. Using Olfaction to Study Memory. Ann. N. Y. Acad. Sci. 855, 657–669 (1998).

6. Gire, D. H., Kapoor, V., Arrighi-Allisan, A., Seminara, A. & Murthy, V. N. Mice develop efficient strategies for foraging and navigation using complex natural stimuli. Curr. Biol. CB 26, 1261–1273 (2016).

7. Yilmaz, M. & Meister, M. Rapid innate defensive responses of mice to looming visual stimuli. Curr. Biol. CB 23, 2011–2015 (2013).

8. Edut, S. & Eilam, D. Rodents in open space adjust their behavioral response to the different risk levels during barn-owl attack. BMC Ecol. 3, 10 (2003).

9. Muenzinger, K. & Gentry, E. Tone discrimination in white rats. J. Comp. Psychol. 12, 195–206 (1931).

10. Tolman, E. C. Prediction of vicarious trial and error by means of the schematic sowbug. Psychol. Rev. 46, 318–336 (1939).

11. Tolman, E. C. Cognitive maps in rats and men. Psychol. Rev. 55, 189–208 (1948).

12. Redish, A. D. Vicarious trial and error. Nat. Rev. Neurosci. 17, 147–159 (2016).

13. Moser, E. I., Kropff, E. & Moser, M.-B. Place Cells, lGrid Cells, and the Brain’s Spatial Representation System. Annu. Rev. Neurosci. 31, 69–89 (2008).

14. Alexander, A. S. & Nitz, D. A. Spatially Periodic Activation Patterns of Retrosplenial Cortex Encode Route Sub-spaces and Distance Traveled. Curr. Biol. 27, 1551–1560.e4 (2017).

15. Carandini, M. & Churchland, A. K. Probing perceptual decisions in rodents. Nat. Neurosci. 16, 824–831 (2013).

16. Odoemene, O., Nguyen, H. & Churchland, A. K. Visual evidence accumulation behavior in unrestrained mice. bioRxiv 195792 (2017). doi:10.1101/19579.

17. Brunton, B. W., Botvinick, M. M. & Brody, C. D. Rats and Humans Can Optimally Accumulate Evidence for Decision-Making. Science 340, 95–98 (2013).

18. Karlsson, M. P., Tervo, D. G. R. & Karpova, A. Y. Network Resets in lMedial Prefrontal Cortex Mark the Onset of Behavioral Uncertainty. Science 338, 135–139 (2012).

19. Dombeck, D. A. & Reiser, M. B. Real neuroscience in virtual worlds. Curr. Opin. Neurobiol. 22, 3–10 (2012).

20. Minderer, M., Harvey, C. D., Donato, F. & Moser, E. I. Neuroscience: Virtual reality exploredtitle>. Nature 533, 324–325 (2016).

21. Domnisoru, C., Kinkhabwala, A. A. & Tank, D. W. Membrane potential dynamics of grid cells. Nature 495, 199–204 (2013).

22. Adesnik, H. Synaptic Mechanisms of Feature Coding in the Visual Cortex of Awake Mice. Neuron 95, 1147–1159.e4 (2017).

23. Olsen, S. R., Bortone, D. S., Adesnik, H. & Scanziani, M. Gain control by layer six in cortical circuits of vision. Nature 483, 47–52 (2012).

24. Dombeck, D. A., Khabbaz, A. N., Collman, F., Adelman, T. L. & Tank, D. W. Imaging large-scale neural activity with cellular resolution in awake, mobile mice. Neuron 56, 43–57 (2007).

25. Kislin, M. et al. Flat-floored Air-lifted Platform: A New Method for Combining Behavior with Microscopy or Electrophysiology on Awake Freely Moving Rodents. J. Vis. Exp. (2014). doi:10.3791/5186.

26. Nashaat, M. A., Oraby, H., Sachdev, R. N. S., Winter, Y. & Larkum, M. E. Air-Track: a real-world floating environment for active sensing in head-fixed mice. J. Neurophysiol. 116, 1542–1553 (2016).

27. Ravassard, P. et al. Multisensory Control of Hippocampal Spatiotemporal Selectivity. Science 340, 1342–1346 (2013).

28. Aghajan, Z. M. et al. Impaired spatial selectivity and intact phase precession in two-dimensional virtual reality. Nat. Neurosci. 18, 121–128 (2015).

29. Yoon, K., Lewallen, S., Kinkhabwala, A. A., Tank, D. W. & Fiete, I. R. Grid Cell Responses in 1D Environments Assessed as Slices through a 2D Lattice. Neuron 89, 1086–1099 (2016).

30. Schmidt-Hieber, C. & Häusser, M. Cellular mechanisms of spatial navigation in the medial entorhinal cortex. Nat. Neurosci. 16, 325–331 (2013).

31. O’Connor, D. H. et al. Vibrissa-Based Object Localization in Head-Fixed Mice. J. Neurosci. 30, 1947–1967 (2010).

32. Huber, D. et al. Multiple dynamic representations in the motor cortex during sensorimotor learning. Nature 484, 473–478 (2012).

33. Guo, Z. V. et al. Procedures for Behavioral Experiments in Head-Fixed Mice. PLOS ONE 9, e88678 (2014).

34. Burgess, C. P. et al. High-Yield Methods for Accurate Two-Alternative Visual Psychophysics in Head-Fixed Mice. Cell Rep. 20, 2513–2524 (2017).

35. Sanders, J. I. & Kepecs, A. Choice ball: a response interface for two-choice psychometric discrimination in head-fixed mice. J. Neurophysiol. 108, 3416–3423 (2012).

36. Goard, M. J., Pho, G. N., Woodson, J. & Sur, M. Distinct roles of visual, parietal, and frontal motor cortices in memory-guided sensorimotor decisions. eLife 5, e13764 (2016).

37. Harvey, C. D., Coen, P. & Tank, D. W. Choice-specific sequences in parietal cortex during a virtual-navigation decision task. Nature 484, 62–68 (2012).

38. Runyan, C. A., Piasini, E., Panzeri, S. & Harvey, C. D. Distinct timescales of population coding across cortex. Nature 548, 92–96 (2017).

39. Scott, B. B. et al. Fronto-parietal Cortical Circuits Encode Accumulated Evidence with a Diversity of Timescales. Neuron 95, 385–398.e5 (2017).

40. Kepecs, A., Uchida, N., Zariwala, H. A. & Mainen, Z. F. Neural correlates, computation and behavioural impact of decision confidence. Nature 455, 227–231 (2008).

41. Uchida, N. & Mainen, Z. F. Speed and accuracy of olfactory discrimination in the rat. Nat. Neurosci. 6, 1224–1229 (2003).

42. Kelemen, E. & Fenton, A. A. Dynamic Grouping of Hippocampal Neural Activity During Cognitive Control of Two Spatial Frames. PLOS Biol. 8, e1000403 (2010).

43. Schmitt, L. I. et al. Thalamic amplification of cortical connectivity sustains attentional control. Nature 545, 219–223 (2017).

44. Towal, R. B. & Hartmann, M. J. Right-left asymmetries in the whisking behavior of rats anticipate head movements. J. Neurosci. 26, 8838–8846 (2006).

45. Aronov, D. & Tank, D. W. Engagement of neural circuits underlying 2D spatial navigation in a rodent virtual reality system. Neuron 84, 442–456 (2014).

46. Dickman, J. D. & Angelaki, D. E. Vestibular convergence patterns in vestibular nuclei neurons of alert primates. J. Neurophysiol. 88, 3518–3533 (2002).

47. Shinder, M. E. & Taube, J. S. Resolving the Active versus Passive Conundrum for Head Direction Cells. Neuroscience 0, 123–138 (2014).

48. Liu, B., Huberman, A. D. & Scanziani, M. Cortico-fugal output from visual cortex promotes plasticity of innate motor behaviour. Nature 538, 383–387 (2016).

49. Knierim, J. J., Kudrimoti, H. S. & McNaughton, B. L. Place cells, head direction cells, and the learning of landmark stability. J. Neurosci. 15, 1648–1659 (1995).

50. Czurkó, A., Hirase, H., Csicsvari, J. & Buzsáki, G. Sustained activation of hippocampal pyramidal cells by ‘space clamping’ in a running wheel. Eur. J. Neurosci. 11, 344–352 (1999).

51. Shapiro, M. L., Tanila, H. & Eichenbaum, H. Cues that hippocampal place cells encode: dynamic and hierarchical representation of local and distal stimuli. Hippocampus 7, 624–642 (1997).

52. Chen, G., King, J. A., Lu, Y., Cacucci, F. & Burgess, N. Spatial cell firing during virtual navigation of open arenas by head-restrained mice. bioRxiv 246744 (2018). doi:10.1101/24674.

53. Cai, D. J. et al. A shared neural ensemble links distinct contextual memories encoded close in time. Nature 534, 115–118 (2016).

54. Ghosh, K. K. et al. Miniaturized integration of a fluorescence microscope. Nat. Methods 8, 871–878 (2011).

55. Fee, M. S. & Leonardo, A. Miniature motorized microdrive and commutator system for chronic neural recording in small animals. J. Neurosci. Methods 112, 83–94 (2001).

56. Voigts, J., Siegle, J. H., Pritchett, D. L. & Moore, C. I. The flexDrive: an ultra-light implant for optical control and highly parallel chronic recording of neuronal ensembles in freely moving mice. Front. Syst. Neurosci. 7, (2013).

57. Buzsáki, G. et al. Tools for probing local circuits: high-density silicon probes combined with optogenetics. Neuron 86, 92–105 (2015).

58. Long, M. A. & Lee, A. K. Intracellular recording in behaving animals. Curr. Opin. Neurobiol. 22, 34–44 (2012).

59. Lee, D. & Lee, A. K. Whole-Cell Recording in the Awake Brain. Cold Spring Harb. Protoc. 2017, pdb.top087304 (2017).

60. Lee, A. K., Manns, I. D., Sakmann, B. & Brecht, M. Whole-Cell Recordings in lFreely Moving Rats. Neuron 51, 399–407 (2006).

61. Epsztein, J., Brecht, M. & Lee, A. K. Intracellular determinants of hippocampal CA1 place and silent cell activity in a novel environment. Neuron 70, 109–120 (2011).

62. Helmchen, F., Fee, M. S., Tank, D. W. & Denk, W. A miniature head-mounted two-photon microscope. high-resolution brain imaging in freely moving animals. Neuron 31, 903–912 (2001).

63. Zong, W. et al. Fast high-resolution miniature two-photon microscopy for brain imaging in freely behaving mice. Nat. Methods 14, 713–719 (2017).

64. Denk, W., Strickler, J. H. & Webb, W. W. Two-photon laser scanning fluorescence microscopy. Science 248, 73–76 (1990).

65. Helmchen, F. & Denk, W. Deep tissue two-photon microscopy. Nat. Methods 2, 932–940 (2005).

66. Theer, P., Hasan, M. T. & Denk, W. Two-photon imaging to a depth of 1000 μm in living brains by use of a Ti:Al_2O_3 regenerative amplifier. Opt. Lett. 28, 1022 (2003).

67. Sofroniew, N. J., Flickinger, D., King, J. & Svoboda, K. A large field of view two-photon mesoscope with subcellular resolution for in vivo imaging. eLife 5, e14472 (2016).

68. Tian, L. et al. Imaging neural activity in worms, flies and mice with improved GCaMP calcium indicators. Nat. Methods 6, 875–881 (2009).

69. Packer, A. M. et al. Two-photon optogenetics of dendritic spines and neural circuits in 3D. Nat. Methods 9, 1202–1205 (2012).

70. Packer, A. M., Russell, L. E., Dalgleish, H. W. P. & Häusser, M. Simultaneous all-optical manipulation and recording of neural circuit activity with cellular resolution in vivo. Nat. Methods 12, 140–146 (2015).

71. Andrasfalvy, B. K., Zemelman, B. V., Tang, J. & Vaziri, A. Two-photon single-cell optogenetic control of neuronal activity by sculpted light. Proc. Natl. Acad. Sci. 107, 11981–11986 (2010).

72. Jun, J. J. et al. Fully integrated silicon probes for high-density recording of neural activity. Nature 551, 232–236 (2017).

73. Hamill, O. P., Marty, A., Neher, E., Sakmann, B. & Sigworth, F. J. Improved patch-clamp techniques for high-resolution current recording from cells and cell-free membrane patches. Pflugers Arch. 391, 85–100 (1981).

74. Bittner, K. C., Milstein, A. D., Grienberger, C., Romani, S. & Magee, J. C. Behavioral time scale synaptic plasticity underlies CA1 place fields. Science 357, 1033–1036 (2017).

75. Harvey, C. D., Collman, F., Dombeck, D. A. & Tank, D. W. Intracellular dynamics of hippocampal place cells during virtual navigation. Nature 461, 941–946 (2009).

76. Kodandaramaiah, S. B., Franzesi, G. T., Chow, B. Y., Boyden, E. S. & Forest, C. R. Automated whole-cell patch-clamp electrophysiology of neurons in vivo. Nat. Methods 9, 585–587 (2012).

77. Kodandaramaiah, S. B. et al. Assembly and operation of the autopatcher for automated intracellular neural recording in vivo. Nat. Protoc. 11, 634–654 (2016).

78. Suk, H.-J. et al. Closed-Loop Real-Time Imaging Enables Fully Automated Cell-Targeted Patch-Clamp Neural Recording In Vivo. Neuron 95, 1037–1047.e11 (2017).

79. Poulet, J. F. A. & Petersen, C. C. H. Internal brain state regulates membrane potential synchrony in barrel cortex of behaving mice. Nature 454, 881–885 (2008).

80. Taube, J. S., Muller, R. U. & Ranck, J. B. Head-direction cells recorded from the postsubiculum in freely moving rats. I. Description and quantitative analysis. J. Neurosci. 10, 420–435 (1990).

81. Colgin, L. L., Moser, E. I. & Moser, M.-B. Understanding memory through hippocampal remapping. Trends Neurosci. 31, 469–477 (2008).

82. Pachitariu, M. et al. Suite2p: beyond 10,000 neurons with standard two-photon microscopy. bioRxiv 061507 (2016). doi:10.1101/06150.

83. Pnevmatikakis, E. A. & Giovannucci, A. NoRMCorre: nAn online algorithm for piecewise rigid motion correction of calcium imaging data. J. Neurosci. Methods 291, 83–94 (2017).

84. Mao, D., Kandler, S., McNaughton, B. L. & Bonin, V. Sparse orthogonal population representation of spatial context in the retrosplenial cortex. Nat. Commun. 8, 243 (2017).

85. Goldey, G. J. et al. Removable cranial windows for long-term imaging in awake mice. Nat. Protoc. 9, 2515–2538 (2014).

86. Kim, T. H. et al. Long-Term Optical Access to an Estimated One Million Neurons in the Live Mouse Cortex. Cell Rep. 17, 3385–3394 (2016).

87. Chen, T.-W. et al. Ultrasensitive fluorescent proteins for imaging neuronal activity. Nature 499, 295–300 (2013).

88. Guizar-Sicairos, M., Thurman, S. T. & Fienup, J. R. Efficient subpixel image registration algorithms. Opt. Lett. 33, 156–158 (2008).

89. Sofroniew, N. J., Vlasov, Y. A., Hires, S. A., Freeman, J. & Svoboda, K. Neural coding in barrel cortex during whisker-guided locomotion. eLife 4,

90. Barretto, R. P. J. et al. Time-lapse imaging of disease progression in deep brain areas using fluorescence microendoscopy. Nat. Med. 17, 223–228 (2011).

91. Mizrahi, A., Crowley, J. C., Shtoyerman, E. & Katz, L. C. High-Resolution In Vivo Imaging of Hippocampal Dendrites and Spines. J. Neurosci. 24, 3147–3151 (2004).

92. Low, R. J., Gu, Y. & Tank, D. W. Cellular resolution optical access to brain regions in fissures: nImaging medial prefrontal cortex and grid cells in entorhinal cortex. Proc. Natl. Acad. Sci. 111, 18739–18744 (2014).

93. Andermann, M. L. et al. Chronic Cellular Imaging of Entire Cortical Columns in Awake Mice Using Microprisms. Neuron 80, 900–913 (2013).

94. Chen, J. L., Pfäffli, O. A., Voigt, F. F., Margolis, D. J. & Helmchen, F. Online correction of licking-induced brain motion during two-photon imaging with a tunable lens. J. Physiol. 591, 4689–4698 (2013).

95. Andermann, M. L., Kerlin, A. M. & Reid, C. Chronic cellular imaging of mouse visual cortex during operant behavior and passive viewing. Front. Cell. Neurosci. 4, (2010).

96. Radvansky, B. A. & Dombeck, D. A. An olfactory virtual reality system for mice. Nat. Commun. 9, 839 (2018).

97. Siegle, J. H., Pritchett, D. L. & Moore, C. I. Gamma-range synchronization of fast-spiking interneurons can enhance detection of tactile stimuli. Nat. Neurosci. 17, 1371–1379 (2014).

98. Rich, P. D., Liaw, H.-P. & Lee, A. K. Large environments reveal the statistical structure governing hippocampal representations. Science 345, 814–817 (2014).

99. Carmein, D. E. E. Omni-directional treadmill. (2000).

100. Carmein, D. E. E. Omni-directional treadmill with applications. (2010).

101. Kaupert, U. et al. Spatial cognition in a virtual reality home-cage extension for freely moving rodents. J. Neurophysiol. 117, 1736–1748 (2017).

102. Lee, K.-S., Huang, X. & Fitzpatrick, D. ON and OFF subfield organization of layer 2/3 neurons in tree shrew visual cortex. J. Vis. 15, 990–990 (2015).

103. Nair, J., Topka, M., Khani, A., Isenschmid, M. & Rainer, G. Tree shrews (Tupaia belangeri) exhibit novelty preference in the novel location memory task with 24-h retention periods. Front. Psychol. 5, 303 (2014).

104. Khani, A. & Rainer, G. Recognition memory in tree shrew (Tupaia belangeri) after repeated familiarization sessions. Behav. Processes 90, 364–371 (2012).

105. Yao, Y.-G. Creating animal models, why not use the Chinese tree shrew (Tupaia belangeri chinensis)? Zool. Res. 38, 118–126 (2017).

106. Newman, J. P. et al. Optogenetic feedback control of neural activity. eLife 4, e07192 (2015).

107. Nashaat, M. A. et al. Pixying Behavior: A Versatile Real-Time and Post-Hoc Automated Optical Tracking Method for Freely Moving and Head Fixed Animals. eNeuro ENEURO.0245-16.2017 (2017). doi:10.1523/ENEURO.0245-16.201.

108. Andermann, M. L., Kerlin, A. M., Roumis, D. K., Glickfeld, L. L. & Reid, R. C. Functional specialization of mouse higher visual cortical areas. Neuron 72, 1025–1039 (2011).

109. Pnevmatikakis, E. A. et al. Simultaneous Denoising, Deconvolution, and Demixing of Calcium Imaging Data. Neuron 89, 285–299 (2016).

